# Wood ants on the edge: how do the characteristics of linear edges effect the population dynamics of an edge specialist?

**DOI:** 10.1101/2025.02.12.637965

**Authors:** Jacob A. Podesta, Catherine L. Parr, Kelly R. Redeker, Elva J. H. Robinson

**Affiliations:** Department of Biology, University of York, York, UK; School of Environmental Sciences, University of Liverpool, Liverpool, UK

**Author notes:** Corresponding author email address (Elva J. H. Robinson).

**Keywords:** Dispersal, Habitat fragmentation, Forest edges, Social insects, Population dynamics, Fire management

## Abstract

Landscape structure modulates species dispersal by presenting barriers or opportunities. Slow-dispersing edge specialists, e.g. the northern hairy wood ant (*Formica lugubris*), are likely to be most affected by topography and land management practices, because they require adjacent contrasting habitats, e.g. to access both food and sunlight. In managed forests, canopy gaps are often linear and anthropogenic, such as paths, firebreaks, and roads (collectively ‘rides’), and their orientation determines shade distribution. Using data spanning 10 years, we ask how ride orientation and width affect the distribution and dispersal of three *F. lugubris* populations in the North York Moors, UK. Ride orientation clearly affected nest abundance, with a higher nest density on rides oriented north-south (N-S) or east-west (E-W) (cardinal directions) than on those oriented NE-SW or NW-SE (intercardinal directions). Conforming to predictions based on sunlight availability, N-S oriented rides were occupied more symmetrically than E-W ones, where the north side was used predominantly.

Nests were generally larger on narrower rides. Ride orientation also clearly affected dispersal: wood ants dispersed c15 m/year along rides oriented in cardinal directions, compared with only c5 m/year on rides oriented intercardinally. Our results show that ride characteristics (width, orientation), resulting directly from forestry practices, influence the distribution and dispersal of an ecosystem engineer woodland species; this may also apply to other forest-edge specialists. As wood ants can suppress defoliating pests, these findings could benefit forest management; forest planners could encourage wood ant colonisation of plantation forest by ensuring linear features contribute to north-south and east-west connectivity.

## Introduction

The ability of species to move through a landscape can have major implications for the conservation of biodiversity, and for preventing species loss in a changing climate (Mendenhall *et al*., 2014; Lenoir and Svenning, 2015). How porous a landscape is to individual movement will also affect the dispersal of species, and this porosity may be determined by the contrast between one habitat and the next. For example, low contrast edges between forest patches and agricultural land in Central American countryside ecosystems are more porous to bats than high contrast edges between forest patches and open water, and forest patches in the agricultural ecosystem harbour higher bat species richness as a result (Mendenhall *et al*., 2014). The permeability of habitats to species may become more important as climate change induced range shifts along latitudinal or altitudinal gradients become more apparent (Lenoir and Svenning, 2015).

Natural linear features such as rivers, coasts, habitat edges, and mountain ranges, or anthropogenic ones such as roads, railways and canals can be both barriers (Shepard *et al.,* 2008) or facilitators of species movement, providing corridors of suitable habitat for certain species through otherwise inhospitable areas. For example, wind-dispersed seeds travel nearly four times the distance along linear forest disturbances than in undisturbed forest (Roberts *et al*., 2018) while, in a meta-analysis, 70 % of studies indicate that either plant abundance or diversity is higher along linear gaps and verges than habitat interiors (Suárez-Esteban *et al*., 2016). By facilitating the dispersal of food plants, linear features can indirectly promote the dispersal of invertebrate herbivores. Ragworts (*Senecio* spp.) disperse along man-made linear features (e.g., railways), and ragwort-dependant cinnabar moths (*Tyria jacobaeae*) show increased dispersal and egg-laying success in areas with roads and valleys compared with areas without linear features (Brunzel et al., 2004), suggesting that invertebrates also benefit from linear features.

The creation of linear features can be a conservation tool, as in the use of habitat corridors to connect smaller areas of protected land and reserves (Bennett, 1990; Brito *et al*., 2017). However, habitat corridors can affect generalist and specialist species differently: landscape corridors connecting areas of fragmented forest in south-eastern Estonia facilitated the dispersal of generalist plant species more than forest specialist species (Liira and Paal, 2013). The presence and structure of edges thus has conservation potential, but the implications for species movement and survival may differ depending on the target species.

The widespread anthropogenic changes in natural forests, and the creation of large areas of plantation forests, mean that human activities determine the present distribution and fragmentation of most forests. This makes the study of how fragmentation and edge habitats impact species dispersal important for conserving species in increasingly anthropogenic habitats. The structure of forest fragments and distribution of edges is especially important in the context of anthropogenic forests because the history of the edges themselves can cause differential effects on different species and their movement. Anthropogenic edges, which are maintained by recent human disturbance, are less permeable to forest specialist carabids (Magura *et al*., 2017) and fungi (Ruete *et al*., 2017) than edges maintained by natural processes. In the context of plantation forest or managed woodland, this highlights the degree of influence that edge creation in forests can have of on species, including edge specialists.

Plantation forests represent a significant portion of forested area in the UK (Defra, 2021) and the management of these forests for timber necessitates canopy gaps (known as rides) to provide access for industrial activity, in addition to firebreaks, geographical features and land ownership boundaries that constrain where trees can be planted. Many of the resulting canopy gaps are linear, here collectively termed ‘rides’. These rides can harbour fauna (Oxbrough *et al.,* 2006) and flora (Smith *et al.,* 2007) distinct from the interior of stands, providing refuges for species that might otherwise be unable to tolerate the plantation environment. Sunlight is an important factor in the differences in species richness and composition between canopy gaps and the interior of stands (Sparks *et al.,* 1996; Smith *et al.,* 2007). The capacity of rides to harbour distinct communities is dependent on ride characteristics, in turn dependent on management practices during ride creation that determine their width and orientation, with downstream effects on community composition (Oxbrough *et al*., 2006; Smith *et al*., 2007). Ride orientation also influences the suitability on rides for certain species and locations: affecting the abundance of ground spiders (Lycosidae) but not ground beetles (Carabidae), for example (Carter *et al*., 1991). While these studies have focused on whole taxonomic groups, we might predict different responses from individual species that are edge specialists.

Certain species are more abundant in edge and transitional habitats between forest and open area because edges allow these species to benefit from two habitats at once. As a result, highly mobile vertebrates able to use multiple habitats, such as the noisy miner (*Manorina melanocephala*; Major *et al*., 2001; Taylor *et al*., 2008; Loyn, 1987) and white-tailed deer (*Odocoileus virginianus*; Williamson and Hirth, 1985), can be edge specialists. Similarly, plants may benefit from improved seed dispersal or pollination opportunities provided by the edge habitat (Lamb and Mallik, 2003) while maintaining the benefits of shelter and nutrient availability (Weathers *et al*., 2001) from nearby forest. Highly mobile species commonly found on edges may merely be ‘edge-tolerant’ generalists exploiting their relative advantage over specialists of the bordering habitats (Ries and Sisk, 2010) rather than having any particular adaptive preference for edges. On the other hand, poor dispersers or sessile species that are more abundant in edge habitats are likely to be dependent on the edge habitat, because of their greater sensitivity to edges (Ewers and Didham, 2005); they will spend multiple generations under its influence and would therefore locally die out if the edge habitat was unfavourable for them. These species may be properly called edge specialists.

The population dynamics of edge specialists are affected by certain properties of edges. For example, the presence of complexities in the forests edge (corners, projections) predict noisy miner (*M. melanocephala*) occupation (Taylor *et al*., 2008). Both edge orientation and width affect butterfly community diversity and composition, with shadier, south-facing edges (southern hemisphere) supporting greater forest butterfly diversity (van Schalkwyk *et al*., 2022) Conversely, leaf miner, parasitoid, and plant communities appear less sensitive to light availability related to edge orientation (Bernaschini *et al*., 2020). While edges clearly impact species distributions, the effect of ride width and especially ride bearing on dispersal are not fully understood.

*Formica lugubris,* a wood ant in the *Formica rufa* group, is a forest edge specialist and a poor disperser in much of its range, including the UK (Ellis and Robinson, 2014). UK populations of *F. lugubris* establish new nests by budding: a mated queen and some workers leave the maternal nest on foot to establish new nests (Maeder *et al*., 2016). As a result, dispersal is local, and even when abundant suitable habitat is available, population margins extend by an average of 5 m per year (Procter *et al*., 2015). These ants are reliant on edge habitat because they require honeydew from arboreal aphids as their main source of carbohydrate but they also require sunlight to regulate the temperature of their nest (Rosengren *et al.,* 1987; Domisch *et al*., 2016). These dual requirements mean that *F. lugubris* nests are typically found within 10m of canopy gaps and on the north side of gaps (Procter *et al*., 2015; Sudd et al., 1977). Their potential for dispersal and population expansion through forest is therefore dependent on the availability of sunlight. In managed forestry land this is largely determined by management decisions regulating the properties of canopy gaps in the form of firebreaks, access roads, footpaths and felled areas.

The orientation of linear canopy openings has the potential to affect both the abundance and dispersal ability of wood ants. In the northern hemisphere, south-facing sites tend to get more insolation. Therefore, along canopy gaps that run approximately east-west, the shadier (south) side of the ride will receive little insolation, so we would predict that wood ants would be less able to utilise nesting sites on the south side of these rides, while canopy gaps that run north-south would be occupied equally on either side, but to a lesser degree, due to the reduced hours in direct sunlight (Figure S.2.). Additionally, wood ant nests found in shadier locations are typically larger; an adaptive response allowing them to thermoregulate more effectively (Chen and Robinson, 2014). As a result, we expect that nests on rides oriented north-south, or nests on the south side of rides running east-west, will be larger than their counterparts on the north side, on average. Finally, we would predict that the width of a ride could affect wood ant abundance: narrow rides where the canopy nearly closes would be shadier, so we would expect larger but less numerous nests on narrow rides and more numerous smaller nests on wider rides.

We conducted a survey of canopy gaps and edge characteristics in plantation forest of the North York Moors at three sites (Cropton Forest, Broxa Forest and East Moor Wood; Figure S.6), and combined pre-existing long-term monitoring data with new data on wood ant populations at the study sites to answer the question: How do characteristics of rides affect the population dynamics of *F. lugubris*? We tested the following hypotheses:

H1 - The orientation of anthropogenic canopy gaps (degrees from N; 0-90°) affects wood ant nest a) abundance and b) average volume of nests along those canopy gaps.
H2 - The orientation of anthropogenic canopy gaps (degrees from N; 0-90°) affects wood ant dispersal rate along those canopy gaps.

## Methods

### Ride selection: abundance and volume measures

Our study site was the North York Moors, UK, where *Formica lugubris* is abundant, populations are expanding into previously unoccupied areas, and high-quality nest distribution data is available from the years 2011, 2013, 2018-2022 (Procter, 2016; Holgate, 2021; Podesta 2023). Rides suitable for testing the hypothesis that canopy gap orientation will affect wood ant nest abundance and volume (H1) were identified prior to the field survey using pre-existing long-term data on wood ant populations in the North York Moors (Procter *et al*., 2015; see Supplementary Methods S.1 and Figure S.6). A ride was defined as a linear feature causing a break in the canopy at least 5m wide, starting and ending where it intersected other rides or reached the end of the forest. Examples of rides included linear canopy gaps due to roads and footpaths, as well as gaps caused by forest management such as areas of row thinning (strips of trees harvested at intervals, leaving most of the crop standing) or firebreaks. We measured the length of the wood ant-occupied portion of the ride from the nest closest to the source population to the nest furthest from it. The population margins and the source were identified using long-term data on the wood ant populations in the North York Moors (Procter, 2016; Holgate, 2021; Podesta, 2023). For rides with nests along their entire length, this measurement equalled the total length of the ride. The total length of all of the rides surveyed was 4133 m. When mapping the margins, a margin was defined as the last occupied area after which no nest was encountered for 200 m; this is far greater than typical dispersal distance for this species, consistent with the long-term data on these populations (Procter *et al*., 2015). Rides were considered to have ended if the ride entered a stand of different tree species, a stand of different age/tree height, or rounded a corner (ride bearing changes by more than 15 degrees). We measured the length and bearing of the ride using the *Measure tool* in ArcGIS Pro version 2.3.3, connecting the first and last nest on the ride.

### Selecting rides for calculating rate of expansion

In order to determine whether ride bearing affects wood ant dispersal, we required a subset of rides at the population margins, where population margin expansion could have occurred during our study period (since 2011) to calculate the average yearly population margins expansion along rides. In addition to the selection criteria in the previous section, margin rides were also required to meet the following criteria:

1. Have had potential for population expansion between 2011 and 2022 i.e., must have uncolonized length that does not extend into unsuitable habitat like open farmland.
2. Expansion has been unidirectional, from a single point of origin, to exclude situations where sections of the ride has been colonised from two origins (due to intersecting two rides with ant presence), as this might inflate the measures of margins expansion on this ride.
3. Expansion has not been interrupted by factors such as areas of forest on the ride being harvested during the period for which we have data, causing the ride to become unsuitable for wood ants. In cases where the expansion had been interrupted, we used the data up to the interruption. For example, a ride that was surveyed from 2011-21, but that was clear-felled in 2019, can provide us with an eight-year average expansion rate rather than ten.

Suitable rides were identified from three forests within the North York Moors, totalling 50 rides, 29 of which were on the margins and suitable for testing margins expansion, each with record of the bearing of the ride, length, mean width, the number of nests on each side of the ride. For the 29 rides on the margins, we also had two measures of the rate of margin change calculated from long-term population data: abundance change (nests/year) and distance change (m/year). One of the rides on the margins was also at the true edge of a forest stand and therefore the ride had no measurable width and only one side on which nests could occur. This ride was dropped from analyses that incorporated side or ride width as variables.

### Nest abundance and volume measurements

For each ride, the surveyor walked down one side of the ride and up the other, recording nest presence and volume. Nests more than 5m from the edge were excluded to ensure all surveyed nests were directly using the ride canopy gap. In areas where the understory was relatively open, spotting nests up to 5m from the edge did not present a challenge but in areas where the understory was dense, some nests were difficult to spot. Where the understory was too thick to confidently spot nests up to 5m from the edge, a second transect was walked, parallel to the first, 2.5 m from the edge (behind obscuring vegetation) to minimise the risk of missing nests. At each nest, we recorded the location (latitude and longitude) and on which side of the ride (east or west/north or south) the nest was, giving each nest a unique ID code to prevent recounting. Additionally, nest width, length and height were measured to calculate nest volume as a proxy for number of ants (Chen and Robinson, 2013).

Ride details were also recorded: these included ride width (the shortest straight line distance between sides of the ride where the canopy is directly overhead and excluding the understory of bushes and seedlings), providing a distance between the crowns of trees, and canopy height (measured using *Toolbox* android app calibrated against a gun clinometer) on both sides of the ride and ride bearing (expressed as degrees east or west of north). We measured each of these ride variables at the beginning, midpoint and end of each ride and averaged them across the three to provide ride characteristics. Latitude and longitude were recorded at the start and end of each ride so that the length of the ride could be measured retrospectively using ArcGIS Pro version 2.3.3.

### Calculating rate of margins expansion along rides

From the long-term data, we calculated two measures of margins change a) distance change and b) number of nests. The distance the margin changed along a ride was the distance in metres between the location of the last nest along a ride recorded in the oldest long-term data and the location of the last nest on the same ride surveyed in 2022. The change in the number of nests was defined as the number of nests between the last nest at the date of the earliest survey in the long-term data and the last nest along the ride in the 2022 survey. Although both rate of distance change and the rate of nest abundance change can be calculated from this data, rate of nest abundance change is a less reliable measure of expansion than the rate of distance change for two reasons. Firstly, there is a five-year interval between measurements in the long-term data and while all population margins were mapped at each survey date, the entire population was not re-mapped on each survey (Procter *et al*., 2015). This means that, in many cases, we have no evidence of continuous occupation, nor of the number of nests that may have been established and subsequently abandoned in the intervening period. Secondly, these populations of *F. lugubris* are polydomous, a life history trait whereby a single ‘colony’ builds and occupies many nest mounds, which share workers and food resources for mutual support. As a result, in areas where the wood ants have recently colonised, some small nests may be abandoned and the occupying ants absorbed into neighbouring nests in the polydomous network (Burns *et al*., 2020) with the effect that the actual numbers of nests may fluctuate. On the other hand, the predicted rate of distance change was low, owing to the slow dispersal of this species (Procter *et al*., 2015) whereas we anticipated that the change in the number of nests would be large and more easily detectable. Note also that the polydomous social organisation of this population means that new nests being established along rides do not necessarily represent new colonies, but rather more likely are an expansion of existing colony networks, although these nests may later become independent.

### Statistical methods

#### Modelling the relationship between ride characteristics and population measures

To test H1 (i.e., the effect of ride characteristics (bearing, width, and side) on the abundance and relative volume of nests) we fitted generalised additive models using the mgcv package version 1.8-41 (Wood, 2011) in R (version 4.2.1). Because our orientation data were circular and we could not make assumptions about the shape of the relationship between orientation and our wood ant population measures, the additional flexibility of a GAM was preferred over a GLM. We fitted models including all combinations of the ride characteristics (bearing, width, and side), including site and ride ID as random effects, and used the minimum AIC to select the most parsimonious model for each response variable (Supplementary Methods S.2). Because of the circular nature of ride bearing (−90° and 90° relative to north are equivalent), we included ride bearing smoothing term as a cyclic p-spline, whereas mean width was included as a p-spline only. We plotted response vs. fitted values, fitted vs. observed and histograms of the residuals to inspect the model fit, and predicted from the models for visualisation. The GAMs provided estimations of degrees of freedom. For nest abundance, cyclic p-splines were fitted for both east and west sides of rides from predicted values of a point count generalised additive model using ride bearing as a predictor offset by ride length, to account for the zero inflation of count data, with the formula nest abundance _≈_ side + s(bearing) + s(width) of the negative binomial family (Link function = log, offset = log(length). For nest volume, the same approach was used, with the formula volume per nest ≈ side + s(bearing) + s(width). For all our models, we calculated deviance explained, as deviance explained is favoured over R^2^ for non-Gaussian models (Table 1, model 1; the nest abundance model) as a more robust measure of explanatory power (Wood, 2006).

**Table 1.**
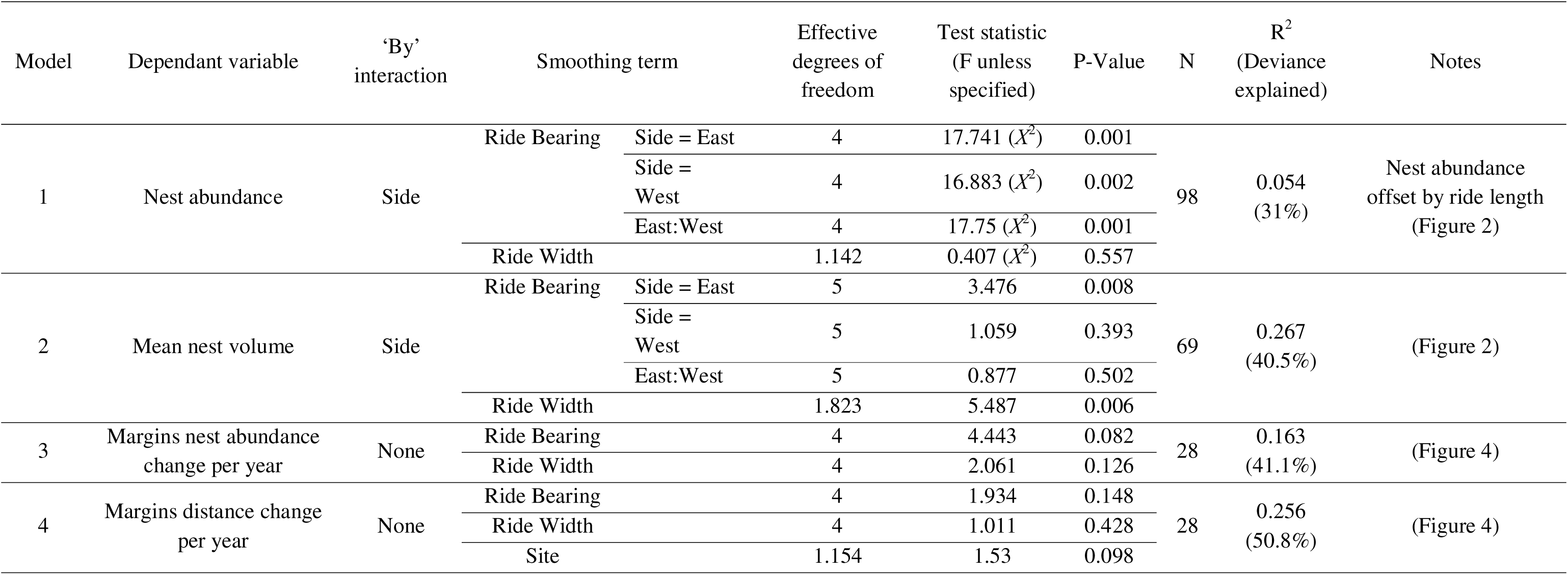
Geralised additive models to predict several wood ant population measures using ride characteristics. Models 1 and 2 included the side of the ride that nests were erm (fitting the two sides as separate splines) so that we could test the significance of each smooths (difference from line with gradient of 0) for data from the ides and the west, and test the smooths against one another (East:West). Both R^2^ and deviance explained are included, as deviance explained is preferred for the n model 1. Expanded table in Supplementary Material (Table S.1.).

#### Pooling bearing into categories for margins growth analysis

Our final dataset resulted in 29 rides available to test H2 (rate of margin change). The bearing data that we collected was continuous, and this was used to fit the GAM model for margins growth as described above. Inspection of the data, however, suggested a bigger difference between rides bearing near the cardinal points (N, S, E, W) and rides bearing near the intercardinals (NE, SW etc.) than between N-S and E-W rides. This suggests that, to test pooled bearing, bins should be E-W +/- 22.5°, N-S +/- 22.5°, and intercardinal +/- 22.5° (Figure S.1.), in order to appropriately capture the variability in the data. This variability has sound ecological justification: assuming that there is a minimum threshold of daily sunlight exposure for nesting sites to be viable for the wood ant nests to persist, we can be sure that this threshold lies somewhere below 50% of summer day lengths, as both sides of N-S rides are often utilised by ants. This threshold is also greater than 0% of the daily hours of sunlight because wood ants were often entirely absent from the south side of E-W rides. As such, going around the compass from north, the hours of available light on the north side will gradually increase from 50% up to nearly 100% after 90°. Meanwhile, the south side will decrease from 50% to nearly 0%, passing below the threshold for wood ants to utilise potential nesting sites somewhere between. This reduction in potential nesting sites and the total availability of sunlight on the ride will have negative effects on wood ant populations and result in slower population growth leading to slower margins expansion.

## Results

### Summary of results

We found that nests were more abundant on the north sides of rides and varied considerably with ride orientation; more nests were found, per metre, on rides that ran E-W or N-S and fewer on rides that were oriented NE-SW or NW-SE (intercardinal). Furthermore, we found that the rate (number of nests) and the distance of dispersal of wood ant nests was lowest along rides intercardinal rides, and three times greater on cardinal rides. On the other hand, no consistent effect was found of ride orientation on nest volume.

### Ride bearing and nest aspect distribution

A total of 371 nests were recorded along all studied rides. Ride orientation covered the full range of possible bearings but rides approximately 15-30° west of north and 70-80° east of north were more abundant than other orientations (Figure 1). The aspect (direction of the long face/lower edge) of nest mounds was overwhelmingly south-facing (Figure 1) with a circular mean of 191.47^°^, significantly different from a uniform distribution (Rayleigh’s test, r = 197.47, var = 0.46, p < 0.0001).

**Figure 1.**
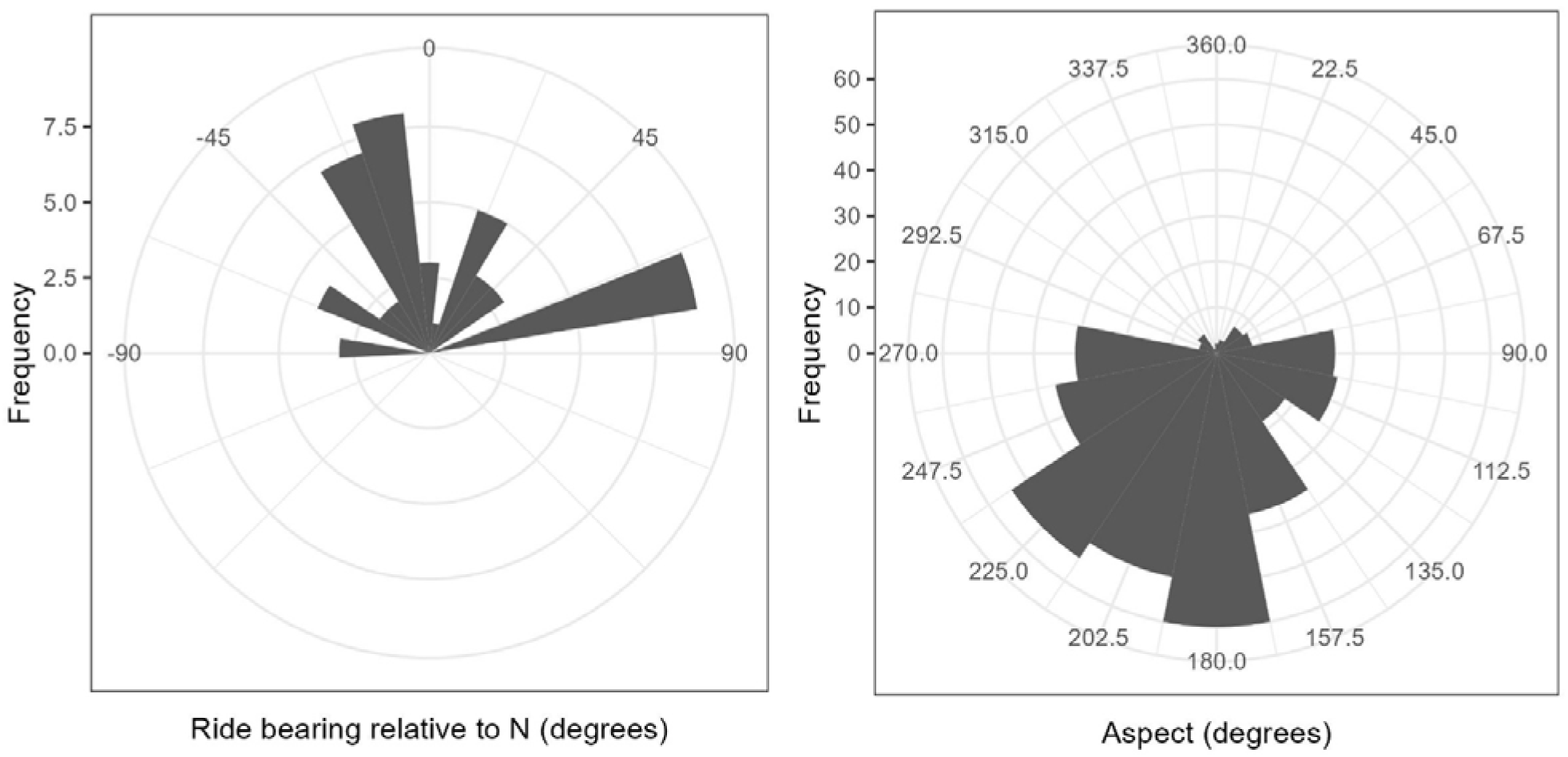
Left: The frequency of rides at different bearings was determined by the availability of suitable rides at the field sites; ride bearings include most possible directions without excessively favouring one direction. Right: The aspect (direction of the lowest side of the nest) of nests was most commonly just west of south (180-225 degrees).

### Does the orientation of canopy gaps affect wood ant nest abundance and volume?

The most parsimonious model of nest abundance included only side, bearing and width (Figure 2; Table 1). There was a significant interaction between ride bearing, and the side of the ride on which nests are most abundant (Figure 2), with nests being most abundant on the East sides of rides that have bearings around the NW-SE axis and most abundant on the West side of rides that have bearings around the NE-SW axis. For rides that have bearings on the NS axis, no difference in abundance between sides was seen. Overall, the number of nests per side was compensated by the other (Figure 2): when the east side has few nests (at bearings that cause the east side to be shaded), the west side has more and vice-versa, so the total sum of nests per ride varied little with bearing. Ride width did not have a significant effect on nest abundance (Table 1).

**Figure 2.**
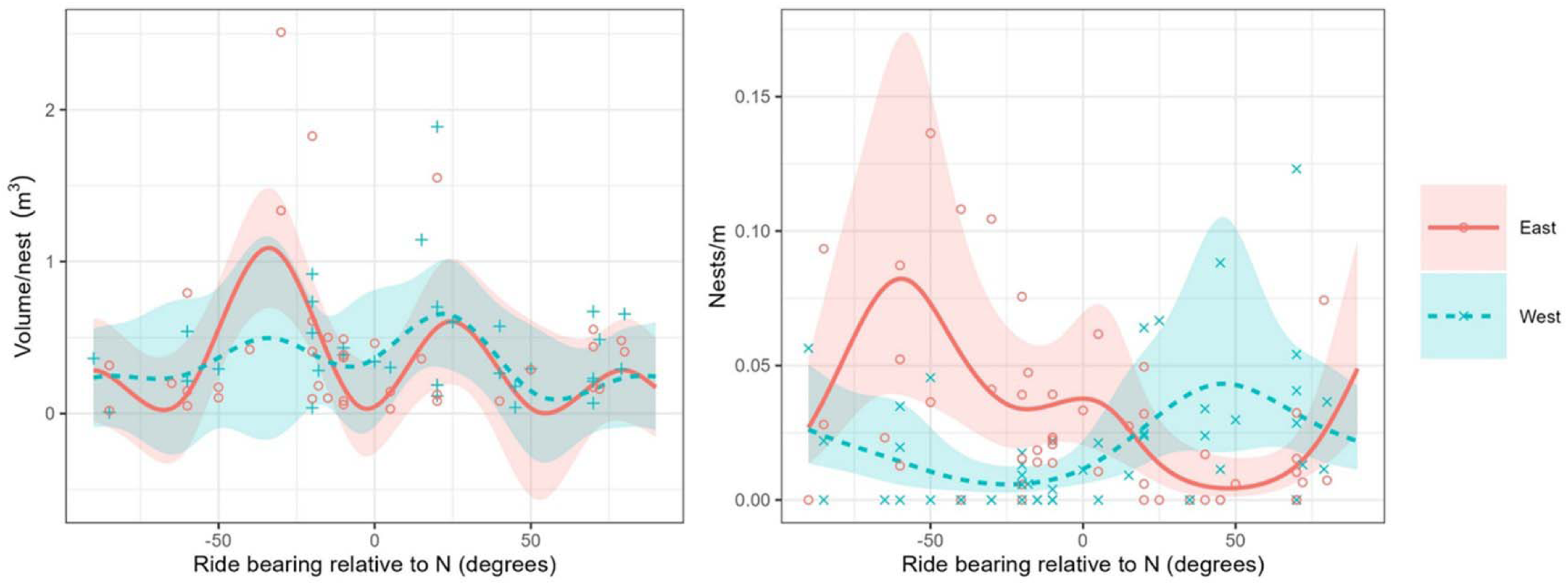
Left: the total volume of all nests on each side of rides of different bearings. The lines of predicted values from a generalised additive model (GAM; R^2^= 0.267, deviance explained = 40.5%) were plotted for values of ride bearing (lines) alongside the original data (points) and 95%CIs (shaded areas). Although the east side smooth was significantly different from a line at the intercept of gradient = 0 (p < 0.01), there was no significant difference between the east and west smooths and no clear pattern is discernible from the plot. Mean width was also significant as a smoothing term (Figure 3; Table 1). Right: the number of nests per side of a ride (E or W) for rides of a range of bearings. The points represent the real data, the lines are predicted values and the shaded areas as are 95%CIs. Both east and west smooths were significantly different from 0 (p < 0.005) and significantly different from each other (p = 0.001).

The most parsimonious model of nest volume included site, bearing and width as smoothing terms, however the inclusion of site only marginally improved explanatory power and did not justify a more complex model, so was dropped (Table S.1; Figure S.3). In both models, the ride bearing significantly affected volume only for nests on the East side, and relationship between ride bearing and nest volume on the East and West sides did not differ significantly from one another (Figure 2; Table 1). The asymmetry of this result, and the lack of difference between the two sides, indicates that overall, ride bearing does not effectively predict mean nest volume in this model. In contrast, mean ride width was significant as a smoothing term (p < 0.01; Table 1) and a plot of model predictions (Figure 3) shows that the mean volume of nests is lower on wider rides but does not continue to decrease as ride width exceeds 8 m. A relatively small number of large nests on rides narrower 6m may be major contributors to this effect; however, this small number of nests were distributed across all three sites.

**Figure 3.**
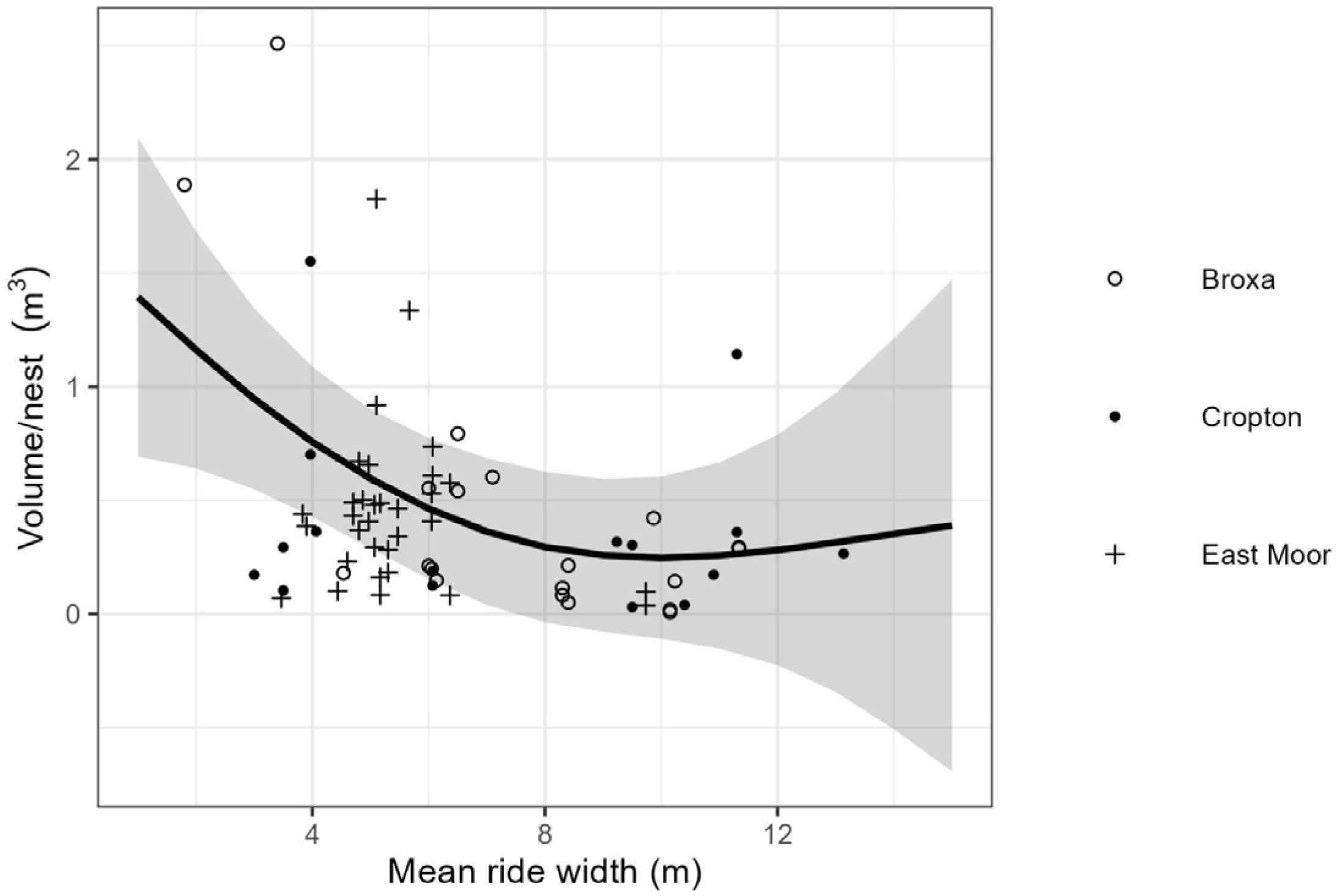
Generalised additive model (GAM) with the formula volume per nest ≈ s(bearing) + s(width), R^2^ = 0.18, deviance explained = 28.7% were plotted for values of width (line) alongside the original data (points) and 95%CIs (shaded area), with the points representing field data. The smooth of mean width was significantly different from the intercept (p < 0.01), whereas ride bearing alone (i.e. not split by side) was not.

### Does the orientation of canopy gaps affect wood ant dispersal rate?

We used two measures of population margins growth: the change in the number of nests at the margin and the distance that the margins moved per year over the study period. The model that best explained the change in nest number per year included ride bearing and width (Figure 4; Table 1). Neither term was below the significance threshold of 0.05 (ride bearing p = 0.082, width p = 0.126) and the model explained only approximately 15% of the variation. Overall, we see a similar shaped pattern in the rate of nest number change (Figure 4) as we saw for nest abundance on each side of rides (Figure 2), with minima for rides bearing approximately NW-SE and NE-SW (intercardinal), though not reaching statistical significance. While this model has poor predictive power, it does suggest that bearing may play some role in the rate of change in nest number.

**Figure 4.**
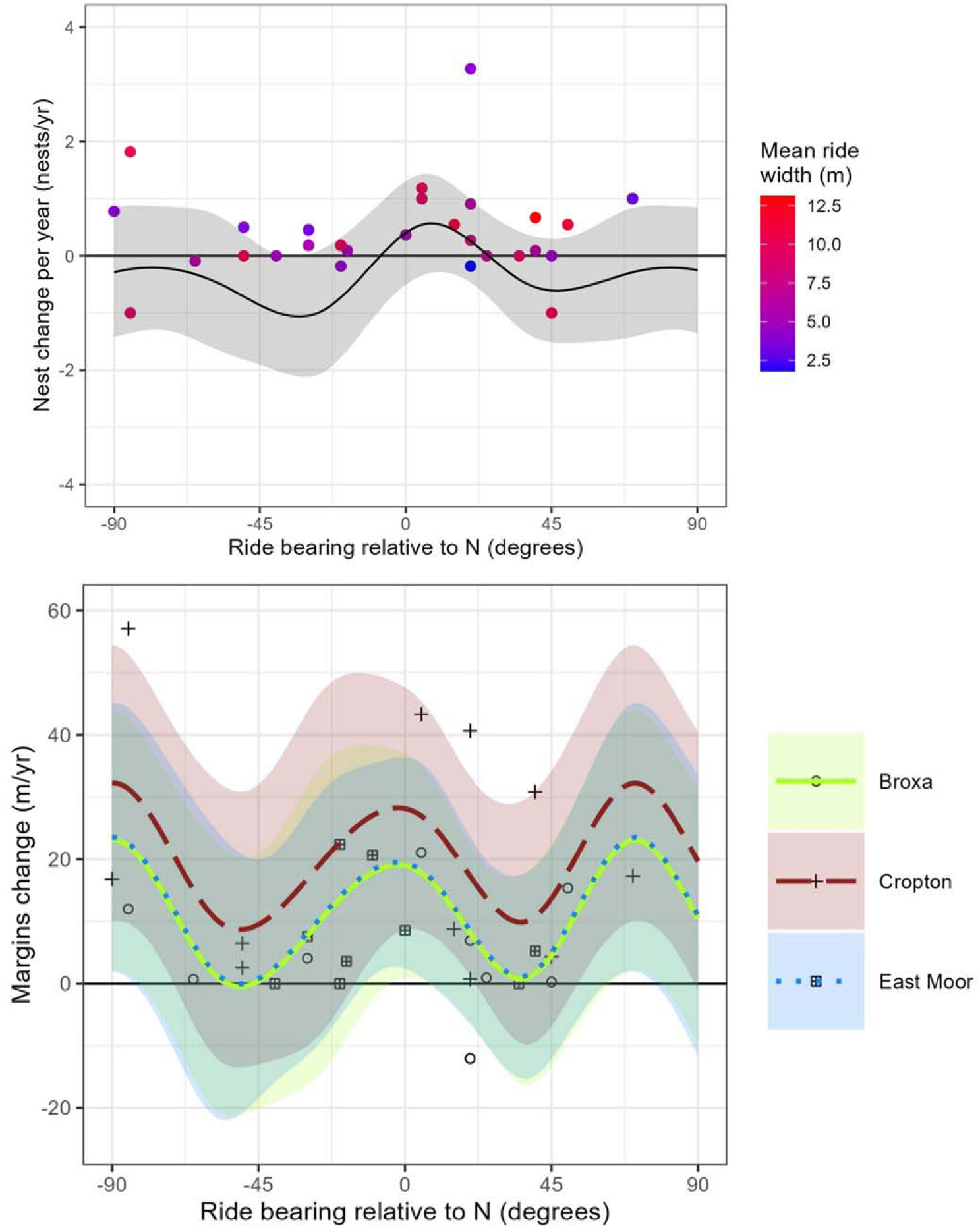
Upper: a generalised additive model (GAM) was fit with the formula nest abundance change per year ≈ s(bearing) + s(width), R^2^ = 0.163, deviance explained = 41.1% and the predictions for values of ride bearing were plotted (solid line) with 95%CIs (shaded areas). Ride bearing was not significant as a smoothing term and the model has little explanatory power. Ride width is not correlated with bearing. Including width as a smoothing term lower predicted values than the observed values due to the fixing of ride width at the mean value for non-Gaussian data. Lower: A generalised additive model was fit with the formula margins change ≈ s(bearing) + s(width) + s(site), R^2^ = 0.256, deviance explained = 50.8%. Empirical data (points), model predictions (lines) and 95%CIs (shaded areas) are shown. None of the smoothing terms were significant, but site was a significant random effect and its inclusion resulted in the most parsimonious model.

The median rate of nest abundance change was greater on rides with cardinal orientations than those with intercardinal ones (approximately 0.5 nests/year; Figure 5), but this difference did not reach statistical significance (Welch’s *t*-test, *t* = 2.02, df = 21.37, p = 0.056). A further split in the data was tested to N-S, E-W, NW-SE and NE-SW against each other, but no statistically significant difference was found between them for either variable (Figures S.4 and S.5).

**Figure 5.**
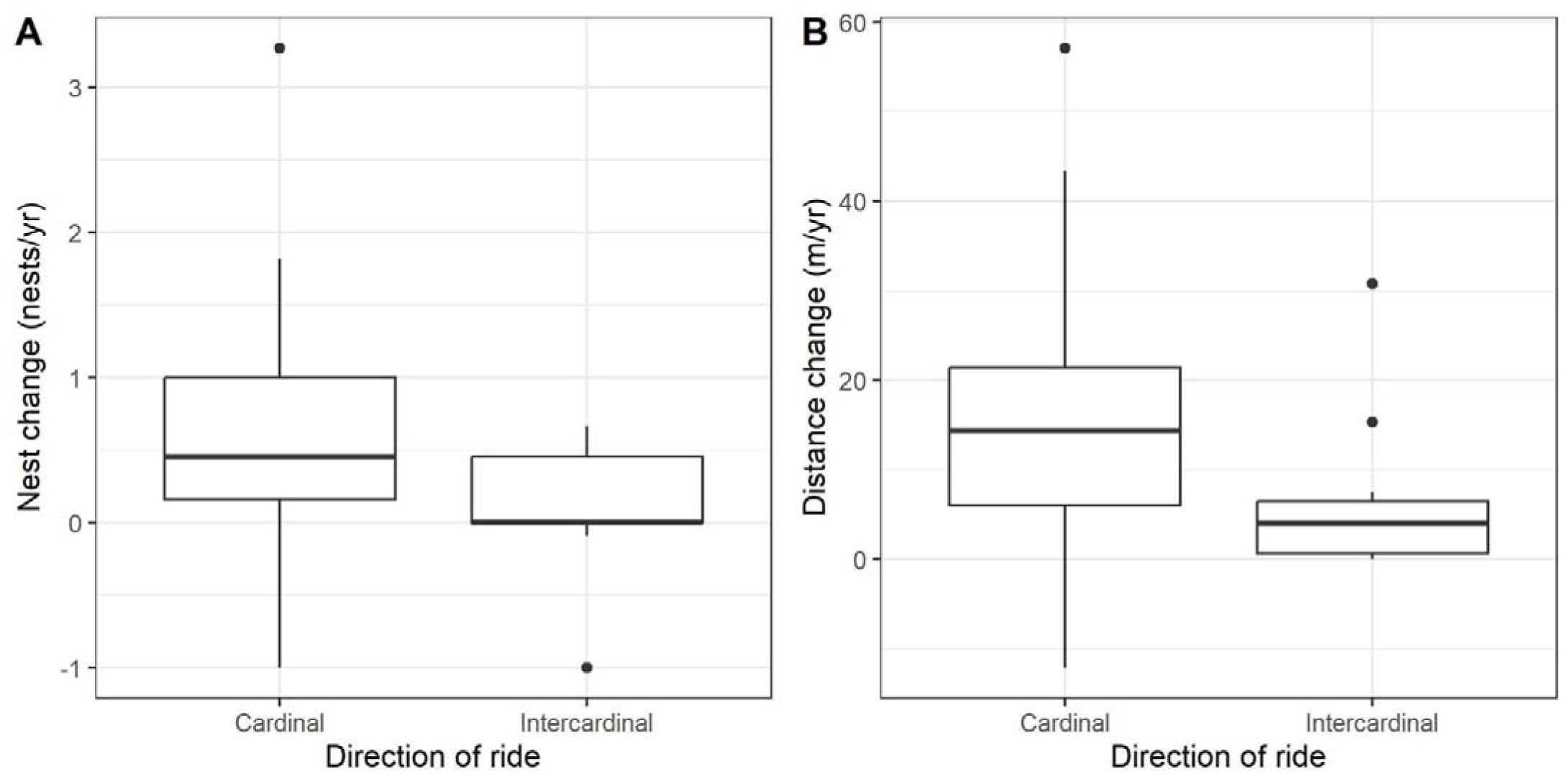
Ride data for change in nest number (A; Welch’s *t*-test, t = 2.02, p = 0.056), and distance of margin change (B; Welch’s *t*-test, t = 2.12, p = 0.045), comparing cardinal (22.5° +/- N, S, E or W) and intercardinal (22.5° +/- NE, NW, SE or SW) bearings (Table 2 for statistical details).

**Table 2.**
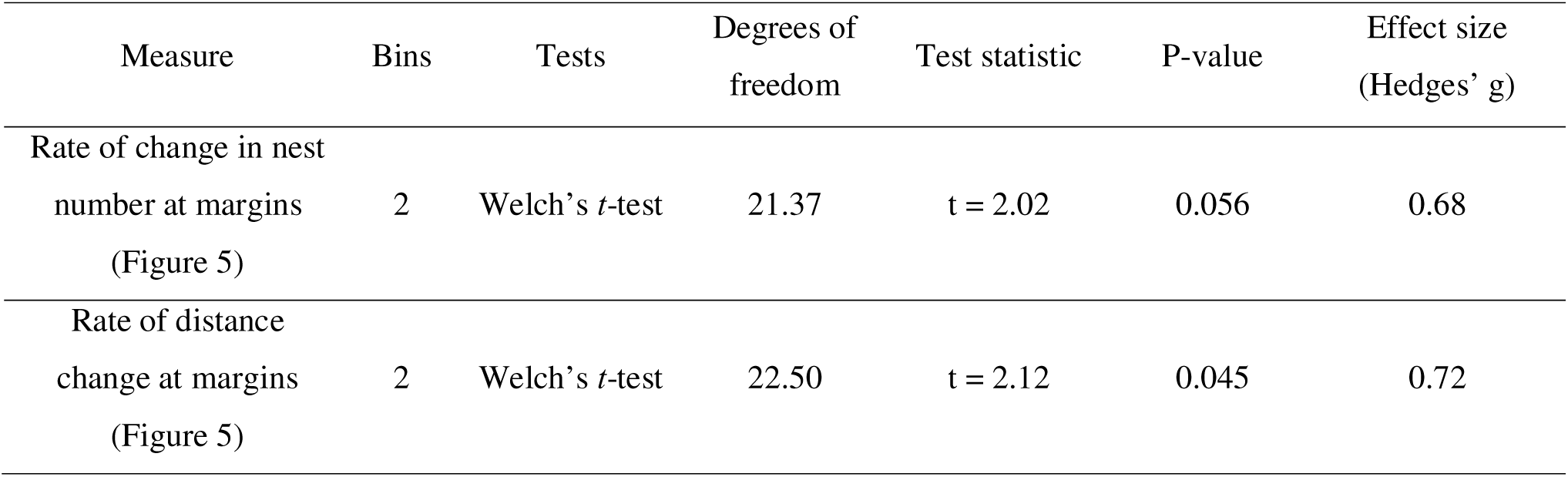
Dispersal range was measured as rate of change of the number of nests at the margin, and distance of the final nest along a ride from the original (2011) population extent to the final (2022) population extent. We made compared rides of varied orientation relative to north; cardinal (N-S and E-W +/- 22.5°) and intercardinal (NW-SE and NE-SW+/- 22.5°) bearings.

| Measure | Bins | Tests | Degrees of freedom | Test statistic | P-value | Effect size (Hedges' g) |
| --- | --- | --- | --- | --- | --- | --- |
| Rate of change in nest number at margins (Figure 5) | 2 | Welch's *t*-test | 21.37 | *t* = 2.02 | 0.056 | 0.68 |
| Rate of distance change at margins (Figure 5) | 2 | Welch's *t*-test | 22.50 | *t* = 2.12 | 0.045 | 0.72 |

The distance that the margins moved per year over the study period was also included as a response variable in a GAM and the best model includes site, ride bearing and ride as smoothing terms (Figure 4; Table 1). This model performed better than the nest number change model (R^2^ = 0.256) and the overall pattern was the same (minima on rides near intercardinal directions), but none of the smoothing terms were significant (p > 0.05; Table 1). Model predictions showed that rides in Cropton Forest experienced more wood ant population growth than either of the other two sites.

Overall, rides running in intercardinal resulted in lower population margins growth (Figure 5). We found that the rides that bear in cardinal directions experienced three times the annual margins expansion than intercardinal ones (15m/year compared to 5m/year; Welch’s *t*-test, *t* = 2.122, df = 22.497, p = 0.045; Figure 5; Table 2) and the effect size was medium (Hedges’ *g* = 0.72).

## Discussion

### Summary of results

We aimed to determine the effect of anthropogenic canopy gap characteristics on populations of an edge specialist, the northern hairy wood ant *Formica lugubris*. Our results clearly show that the orientation and width of linear canopy gaps affect *F. lugubris* distribution and dispersal, shaping their population dynamics. Nests overwhelmingly had a south facing aspect, as would be expected if nests are constructed to present a large surface area to the sun and maximise insolation. Nests were more abundant on the northern side of rides for all orientations, as predicted, but orientation did not similarly affect nest size in a consistent way. In general, narrower rides had larger nests, while wider rides were populated by smaller nests, consistent with predictions that nests receiving less insolation are larger (Chen and Robinson 2014). Finally, we found that, as predicted, the orientation of rides affected *F. lugubris* dispersal. Over the preceding 10 years, populations of *F. lugubris* were both more numerous and dispersed more quickly along rides that run in cardinal directions (N, E, S, W) than ones that run in intercardinal directions (NE, SE, SW, NW).

### The importance of edge characteristics

We showed that the orientation of edge habitats relative to north is an important factor in edge quality for *Formica lugubris* and is sufficient to have measurable effect on nest abundance and population margin expansion. Ride width is also a factor in edge quality for these wood ants; we found larger nests in narrower rides. In general, larger nests are found in shadier areas because larger nests are better able to thermoregulate (Chen and Robinson, 2014). The strong south-facing bias in nest aspect we found confirms that our other results are likely to driven by insolation, as sunlight is important for many aspects of wood ant biology. Nests lacking direct sunlight, especially small nests, may initiate foraging later (Rosengren *et al*., 1987), have slower brood maturation (Kadochová and Frouz, 2013) and therefore potentially slower colony growth.

### Dispersal along rides

In addition to ride bearing affecting wood ant population density (nest volume and abundance), it also affects the ability of the wood ants to disperse through these plantation forests, with most rapid dispersal along rides oriented along the cardinal bearings (N-S or E-W). Due to the generally slow dispersal of *F. lugubris*, the cardinal ride dispersal distance of 15 m/year is equivalent to three times the rate of dispersal in intercardinal rides (5 m/year). We hypothesise that the slower population margin expansion observed for rides with bearings in intercardinal directions is due to a minimum threshold of insolation for newly established nests of *F. lugubris* to be viable. Above this threshold, a wood ant nest’s outcomes continue to improve. We suggest that the south side of a ride with an intercardinal bearing receives insolation below this minimum threshold, due to being shaded for most of the day by the trees on the southern side, while the north side will receive less insolation than the north side of a ride running east-west. This means nests on these bearings are in a ‘worst of both worlds’ environment resulting in fewer individuals and a reduced ability to form new nests in the polydomous network. Polydomy allows for food and labour sharing between nests that can subsidise a new nest while it grows big enough to establish foraging (Ellis and Robinson, 2015). Newly established nests in shadier areas may require extra support from neighbouring nests (Rosengren *et al*., 1987), so being further from the centre of this network may leave new nests more vulnerable to abandonment. The combination of lower rates of nest foundation and a higher failure rate for nests that have been established further from the parental nests, could explain the reduced margins growth that we measured in shadier, intercardinal rides.

### Management implications

Given that wood ants increase biodiversity in plantation forests and may provide additional pest control benefits (Laine and Niemela, 1980; Karhu and Neuvonen 1998; Zingg *et al*., 2018), the results that we present here may be useful in planning management of plantations. Wood ants can reduce defoliating pest burden (Karhu and Neuvonen 1998), influence tree growth via their effects on aphids (Kilpeläinen *et al*., 2009), compete with other insectivores (Haemig, 1992), and change the spatial distribution of soil resources (Podesta, 2023), so the effects of ride characteristics on their dispersal have major implications for conservation and forest management. When planning new areas of forestry adjacent to existing populations of wood ants, or in areas where translocation of wood ants’ nests is being considered (Nielsen *et al*., 2018), we would recommend cutting firebreaks, access tracks etc. in cardinal directions, to maximise the ability of wood ants to disperse. The width of the ride should also be considered when managing forest to promote species richness or dispersal. Previous work has suggested that canopy gaps should be at least 15 m wide in order to harbour species that would be shaded out in the forest interior (Oxbrough *et al.,* 2006; Smith *et al.,* 2007), and our data suggest that wood ants similarly have a minimum threshold of insolation (also impacted by ride width) that allows them to thrive and disperse.

### Wider ecological implications

The work presented here adds to a body of research showing that linear canopy gaps in plantation forests have important implications for forest dwelling species. The characteristics of rides, such as width, affect the diversity or abundance of butterflies (Greatorex-Davies *et al*., 1993), spiders (Oxbrough *et al.,* 2006; Carter *et al.,* 1991), ground beetles (Carter *et al.,* 1991), vascular plants (Sparks *et al.,* 1996; Smith *et al.,* 2007), and bryophytes (Smith *et al.,* 2007). We have shown that, in addition to the effects that ride characteristics can have on the diversity and abundance of forest species in plantation, the orientation of rides can influence the dispersal ability of an edge specialist, opening up the question of whether other species with similar habitat requirements might be similarly affected.

Managing plantations in a manner favourable to wood ant dispersal (by cutting firebreaks and other necessary linear gaps along cardinal directions) could be beneficial to other species and biodiversity in economically productive forests in two ways. Firstly, the presence of wood ants themselves can be beneficial to other species. The nests of wood ants are home to many myrmecophilous species, with over 100 myrmecophile species known from wood ants’ nests (Robinson *et al*., 2016; Parmentier *et al.,* 2014). Additionally, wood ants can play an important role in the dispersal of seed, including some declining and charismatic wildflowers like the endangered *Melampyrum cristatum* (crested cow-wheat) which is dispersed by wood ants and is likely range-limited by the availability of dispersers (Stachnowicz, 2013). Secondly, other species with similar habitat requirements and dispersal ability to the wood ants would benefit from the same rides as dispersal corridors as the wood ants. Species that might benefit from the same corridor design as wood ants could include wildflowers that likewise require direct sunlight but benefit from other conditions provided by the forest edge such as protection from wind, acidic soil or edge related nutrient influx (Weathers *et al*., 2001).

On the other hand, species movement can be detrimental; there are well documented negative economic (Cuthbert *et al*., 2022) and environmental (Pyšek *et al*., 2020) effects of species invasions. Edge specialists with the potential to benefit from improved dispersal due to edge orientation may also be pests, for example, the larvae of *Gilpinia virens,* a pine sawfly and common pest of *Pinus sylvestris*, are found in higher numbers in stands with relatively open canopies (Gawęda and Grodzki, 2020). As a result, pests such as *G. virens* may also benefit from the presence of canopy gaps cut in favourable, cardinal directions, although it should be noted that it does not necessarily follow that a preference for canopy gaps will result in the same response to different gap characteristics as we see in wood ants.

### Conclusions

Our work clearly shows the potential of anthropogenic linear canopy gaps for promoting dispersal in plantation forests. Using a species that favours woodland edges to allow access to both light and resources, we demonstrate that the properties of linear canopy gaps influence distribution and dispersal rates. In practical terms, this means that canopy gaps created during plantation forest management, such as firebreaks, row-thinning, or access for logging machinery, can have the secondary function of promoting the dispersal of edge specialist forest species, especially if orientations near N-S or E-W are predominant. The control that forestry planners have over the placement of firebreaks and other linear canopy gaps, thus allows the possibility of deliberately making plantation forests more porous to certain species, to gain the ecosystem services they provide (for example, the suppression of defoliating pests by wood ants) or to promote biodiversity. The increase in demand for timber products and forested areas to reach carbon offset goals will likely lead to an increase in the area and intensity of plantation forests in the future (McEwan et al., 2020). If these economic and environmental goals to be achieved without detriment to biodiversity or resilience to pests, care must be taken to ensure the development of forestry practices that allow wildlife to proliferate in plantation forests, both for the ecosystem services they can provide and for the biodiversity that they represent.

## Acknowledgements

We thank Duncan Procter and Megan Holgate for additional data that contributed to the work and Forest Research for site access permission. We also thank Imre Sándor Piross and Mark Hodson for advice with the analysis and manuscript respectively, and Anne Oxbrough for feedback on the original thesis chapter. The project was funded by the NERC ACCE2 DTP.

