## Supplementary Material for "Wood ants on the edge: how do the characteristics of linear edges effect the population dynamics of an edge specialist?"

Supplementary Information

Supplementary Methods

S.1 Details of pilot study and power analysis

An initial 13 suitable rides were identified from long-term data at Cropton and Broxa forests (Procter *et al.*, 2015) to assess the feasibility of this study. A power analysis on these data (power = 0.8, alpha = 0.05) suggested that a final sample size of 37 rides would be suitable to test H1 and 16 would be adequate for H2. Based on our desk survey using the long-term data for Cropton and Broxa (Procter *et al.*, 2015), this exceeded the number of rides suitable for each hypothesis at these sites. As a result, additional rides from East Moor Wood in the North York Moors were included. The final number of rides selected for the study was 50, including all 29 of margins rides that were suitable for H2 (site = number of rides/number of rides on the margin; Cropton = 11/11; Broxa = 14/9; East Moor Wood = 25/9). The 21 non-margins rides were selected randomly from Broxa and East Moor Wood.

Our initial power analysis used data gathered from the known nest locations along rides where the population margins were expanding and were close to E-W or N-S. This indicated that 30 rides would be adequate only for an alpha of 0.1, so we aimed to collect more than that. However, this power analysis cut the data such that:

1.	Bearing was categorical.

2.	The difference between N-S and E-W rides was tested.

Suitable rides were identified from three forests within the North York Moors, totalling 50 rides; power analysis on the pre-existing data indicated this would provide sufficient power to test both our hypotheses (power >0.8, alpha=0.05). Ultimately, the resulting data set included 50 rides, 29 of which were on the margins and suitable for testing margins expansion.

One ride is recorded as 90 degrees, which would make recording a nest as on the east or west side of the ride unclear. However, this was due to compass error and inspection of the satellite imagery and maps reveals that the recording one side as eat or west, as we do in our analyses, is reasonable.

S.2. Model selection details

Two models were constructed to predict nest volume: volume per nest ≈ side + s(bearing) + s(width) and volume per nest ≈ side + s(bearing) + s(width) +s(site) (Table S.1; Figure S.3), however the ΔAIC between models with and without site (Table S.1) as a term is 2.278 and only just outside of the widely used cut-off 2 for supporting a more complex model. Furthermore, site was not significant as a (random effect) smoothing term and, despite the slight difference in the mean nest volume of the three sites; its inclusion is unhelpful to the interpretation of the overall pattern and results in a very small increase in R2 (0.267 without site; 0.299 with), given that to do so would inhibit comparison between models.

| **Table S.1.** Generalised additive models to predict several wood ant population measures using ride characteristics. Models 1-3 included the side of the ride that nests were on as a ‘by’ term (fitting the two sides as separate splines) so that we could test the significance of each smooths (difference from line with gradient of 0) for data from the east side of rides and the west, and test the smooths against one another (East:West). Both R^2^ and deviance explained are included, as deviance explained is preferred for the non-Gaussian model 1. AIC is included for the two nest volume models, as ΔAIC was very close to the cut off for accepting the more complex model. | | | | | | | | | | |
| --- | --- | --- | --- | --- | --- | --- | --- | --- | --- | --- |
| Model | Dependant variable | ‘By’ interaction | Smoothing term | | Effective degrees of freedom | Test statistic (F unless specified) | P-Value | N | R^2^  (Deviance explained) | Notes |
| 1 | Nest abundance | Side | Ride Bearing | Side = East | 4 | 17.741 (*X*^2^) | 0.001 | 98 | 0.054  (31%) | Nest abundance  offset by ride length  (Figure 2) |
|  |  |  |  | Side = West | 4 | 16.883 (*X*^2^) | 0.002 |  |  |  |
|  |  |  |  | East:West | 4 | 17.75 (*X*^2^) | 0.001 |  |  |  |
|  |  |  | Ride Width |  | 1.142 | 0.407 (*X*^2^) | 0.557 |  |  |  |
| 2 | Mean nest volume | Side | Ride Bearing | Side = East | 5 | 3.476 | 0.008 | 69 | 0.267  (40.5%) | AIC = 82.64  (Figure 2) |
|  |  |  |  | Side = West | 5 | 1.059 | 0.393 |  |  |  |
|  |  |  |  | East:West | 5 | 0.877 | 0.502 |  |  |  |
|  |  |  | Ride Width |  | 1.823 | 5.487 | 0.006 |  |  |  |
| 3 | Mean nest volume | Side | Ride Bearing | Side = East | 5 | 3.775 | 0.005 | 69 | 0.299  (44.2%) | AIC = 80.36  (Figure S.3) |
|  |  |  |  | Side = West | 5 | 1.157 | 0.342 |  |  |  |
|  |  |  |  | East:West | 5 | 0.877 | 0.502 |  |  |  |
|  |  |  | Ride Width |  | 1.921 | 6.258 | 0.003 |  |  |  |
|  |  |  | Site |  | 0.98 | 1.11 | 0.107 |  |  |  |
| 4 | Mean nest volume | None | Ride Bearing |  | 4 | 1.187 | 0.332 | 46 | 0.18  (28.7%) | (Figure 3) |
|  |  |  | Ride Width |  | 2.164 | 5.053 | 0.009 |  |  |  |
| 5 | Margins nest abundance change per year | None | Ride Bearing |  | 4 | 4.443 | 0.082 | 28 | 0.163  (41.1%) | (Figure 4) |
|  |  |  | Ride Width |  | 4 | 2.061 | 0.126 |  |  |  |
| 6 | Margins distance change per year | None | Ride Bearing |  | 4 | 1.934 | 0.148 | 28 | 0.256  (50.8%) | (Figure 4) |
|  |  |  | Ride Width |  | 4 | 1.011 | 0.428 |  |  |  |
|  |  |  | Site |  | 1.154 | 1.53 | 0.098 |  |  |  |

**Figure S.1.** We pooled the data for margins change for rides of different bearing into two categories for rides running along cardinal and intercardinal directions (+/- 22.50°) so that we could test the differences between more or less exclusive groups respectively.
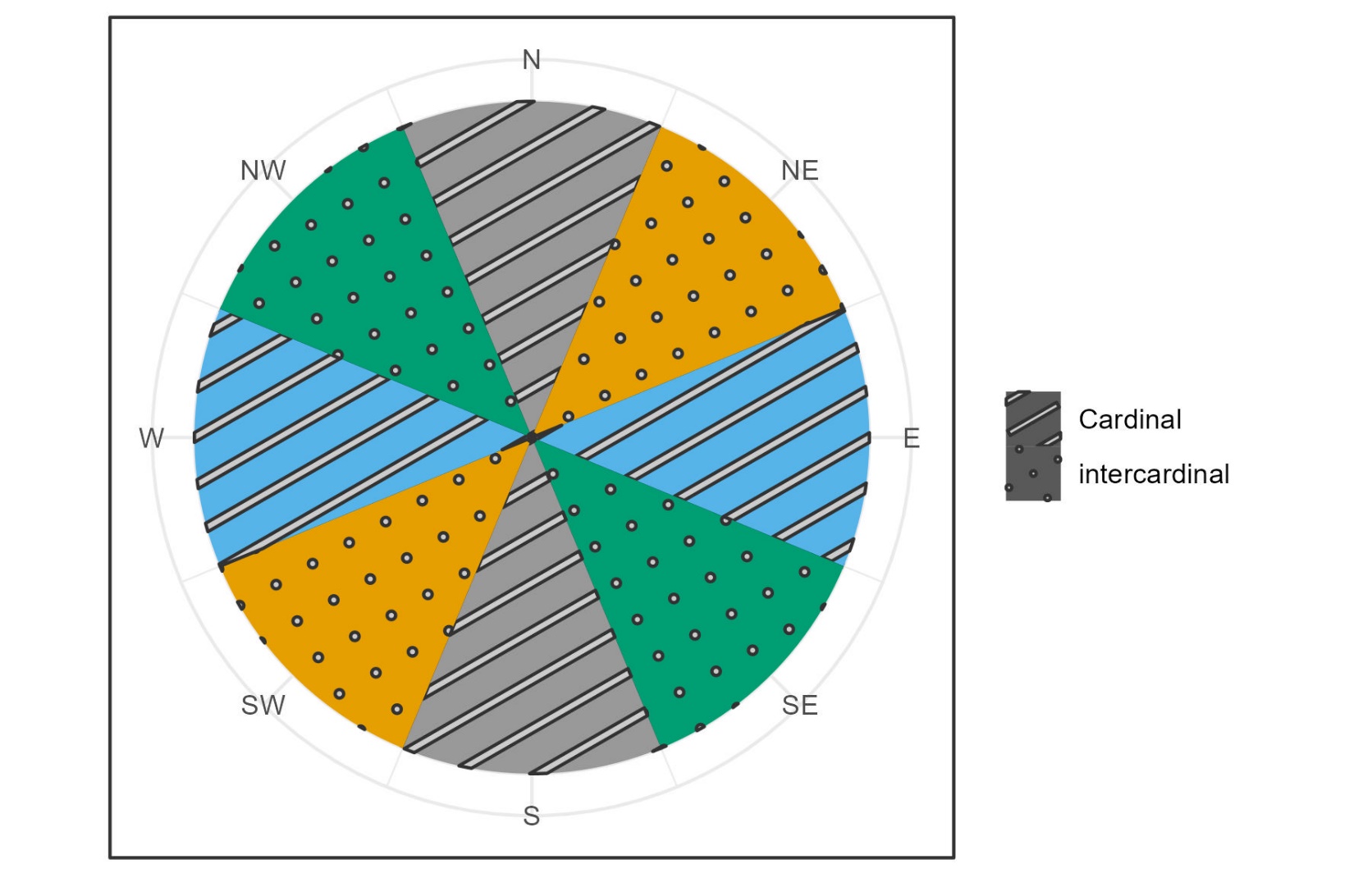

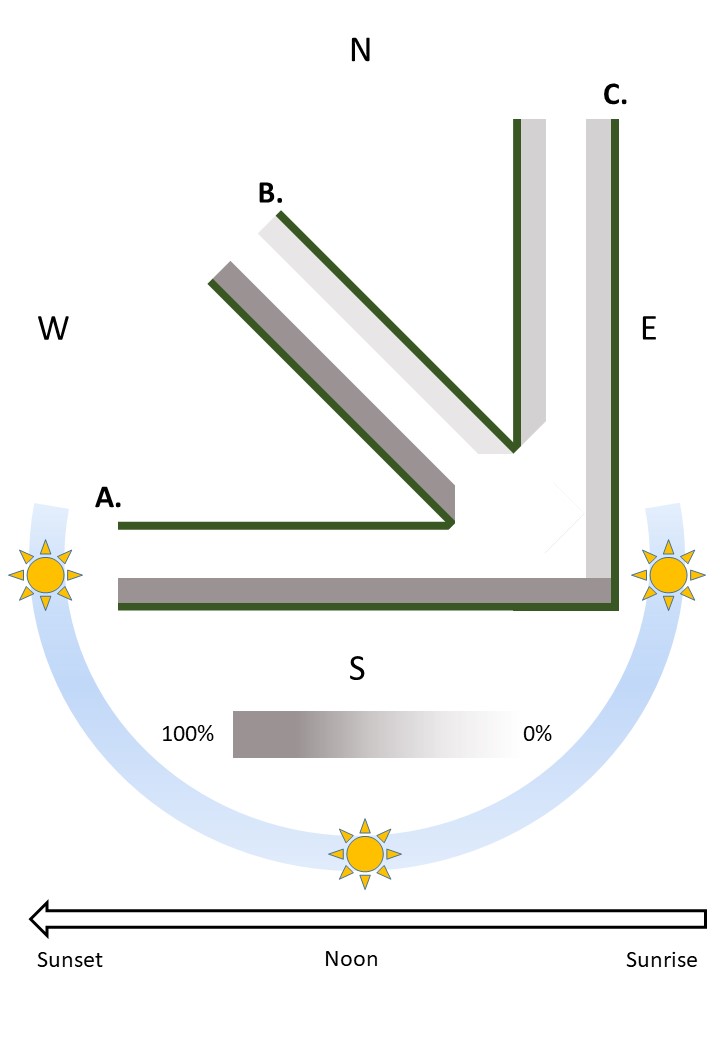
**Figure S.2.** An E-W ride (A) receives a full day of sun on the north side, whereas a N-S ride (C) receives half of a day of sunlight on each side (in the morning on the west side, in the afternoon on the east). Rides in intercardinal directions (e.g., B) will receive fewer hours of sunlight on the north side than and E-W ride, but still receive little or no direct sunlight on the south side. The hours of sunlight available on each side will approach 50% as the ride bearing approaches 0°, while hours of sunlight available will reach maximum asymmetry for rides bearing 90/270° (E-W). Only the north side of the ride A will be suitable for wood ants to nest, and the south will remain unsuitable for all potential bearings until the hours of sunlight reaches a threshold (between B and C) at which both sides can be utilised by the ants but are both sub-optimal compared to the north side of an E-W ride.

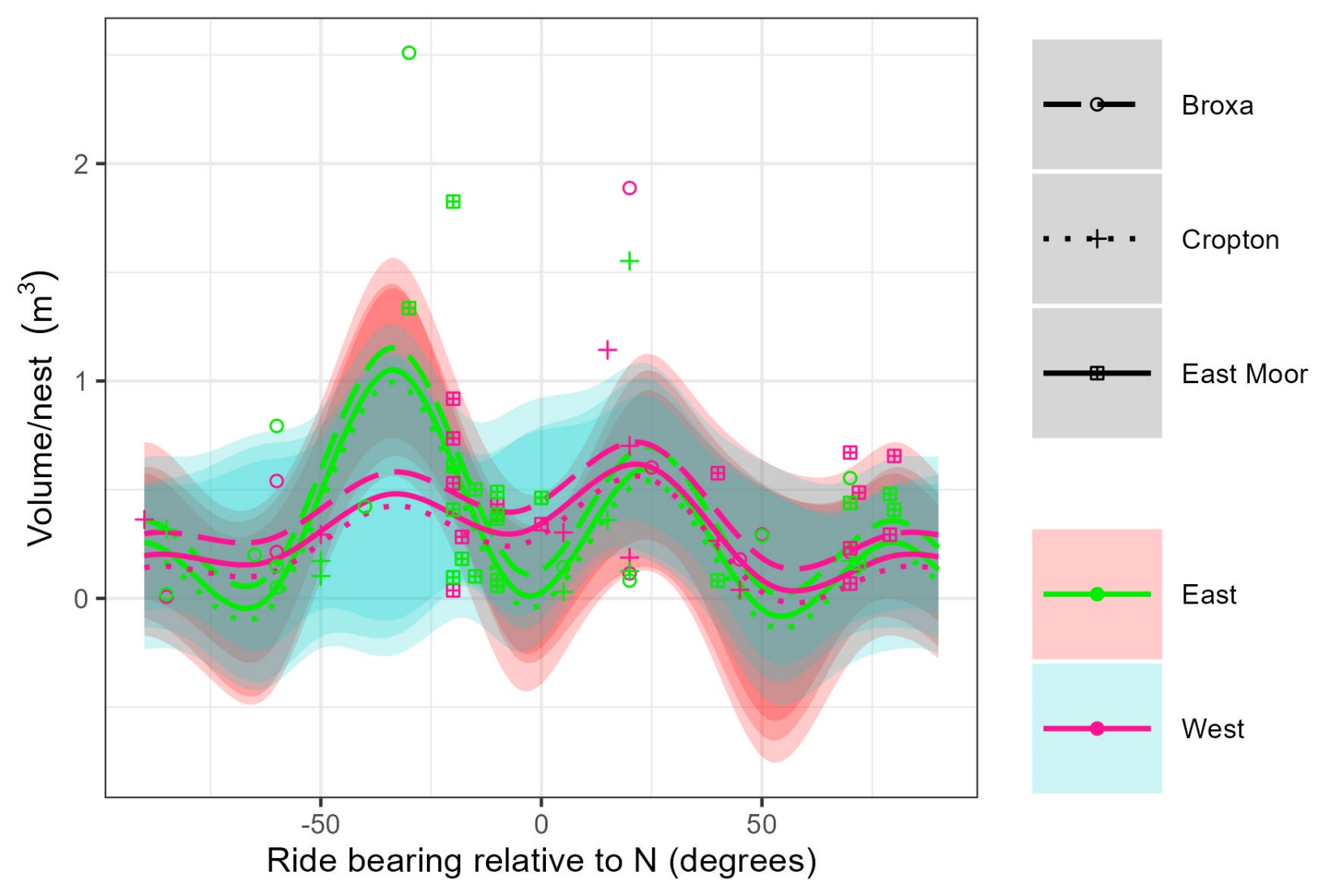
**Figure S.3.** Predicted values from a generalised additive model (GAM) with the formula volume per nest ≈ side + s(bearing) + s(width) + s(site) R^2^ = 0.299, deviance explained = 44.2% were plotted for values of ride bearing (lines) alongside the original data (points) and 95%CIs (shaded areas). Although the east side and mean width smooths were significantly different from a line at the intercept of gradient = 0 (p < 0.01), there was no significant difference between the east and west smooths and no clear pattern is discernible from the plot. Based on AIC values, this was more parsimonious than the model excluding site as a smoothing term (Figure 2), however the ΔAIC = 2.278, only just outside of the widely used cut-off for supporting a more complex model and both models have been included in the supplementary material (Table S.1).

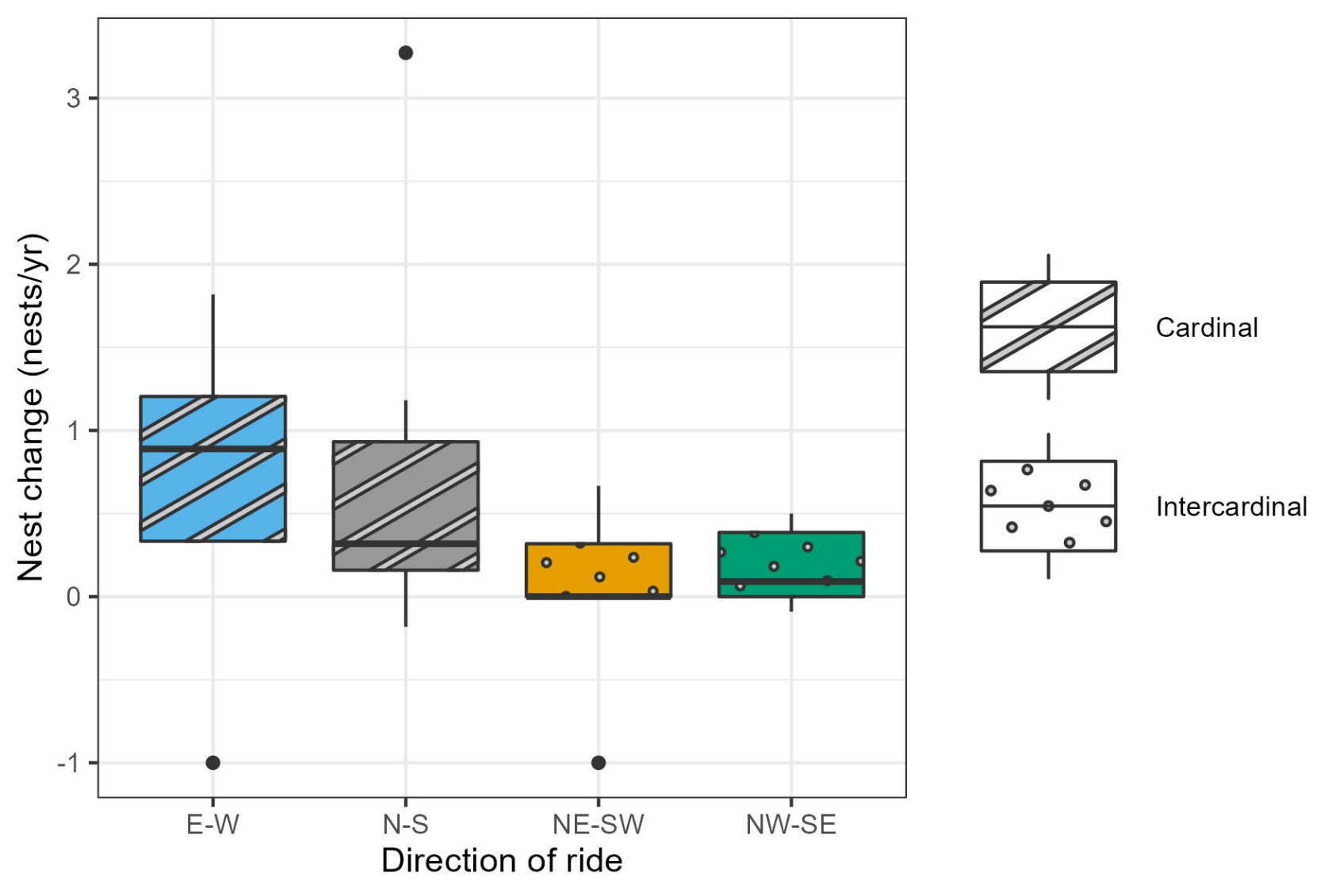
**Figure S.4.** The ride data were pooled by bearing so that the categories consisted of rides bearing +/- 22.5° of the cardinal and intercardinal points. E-W and N-S rides appear to have undergone slightly more (positive) change in nest number than NE-SW and NW-SE rides.

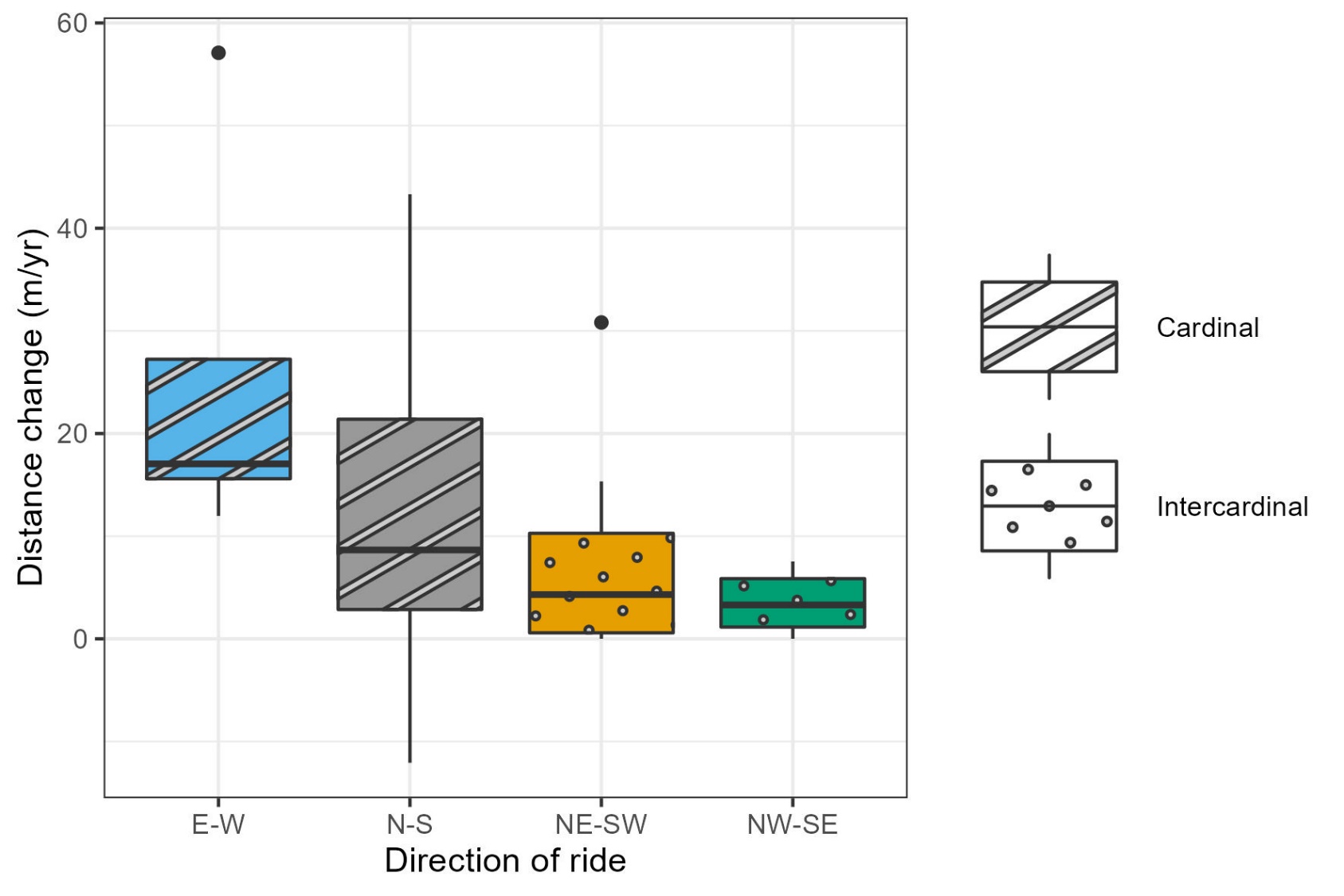
**Figure S.5**. We pooled the data by bearing so that the categories consisted of rides bearing +/- 22.5° of the cardinal and intercardinal points. Although the differences in distance change between category were not significant (ANOVA, p > 0.05), E-W and N-S rides appear to have undergone slightly more (positive) change in margins position than NE-SW and NW-SE rides.

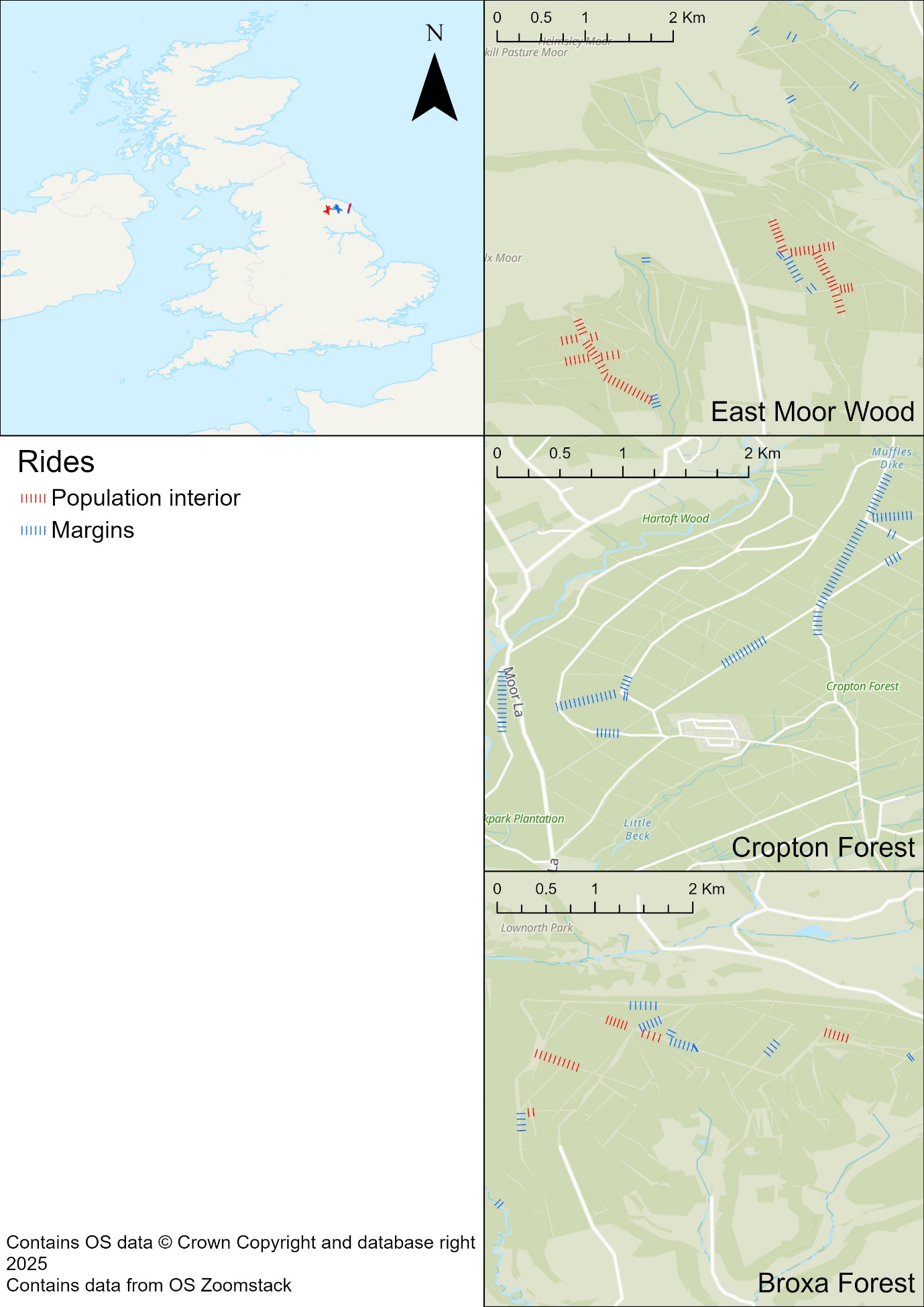

**Figure S.6.** The rides used in this study fell into two categories: population interior, where the whole ride was within the population extent at our earliest data, so population margins expansion into unoccupied areas could not occur at either end of the ride, and margins, where there was unoccupied habitat at least one end of the ride at the date of the earliest data in our study.
